## Supplementary figure 1 for "Variational Bayesian Phylogenies through Matrix Representation of Tree Space"

### SUPPLEMENTARY INFORMATION

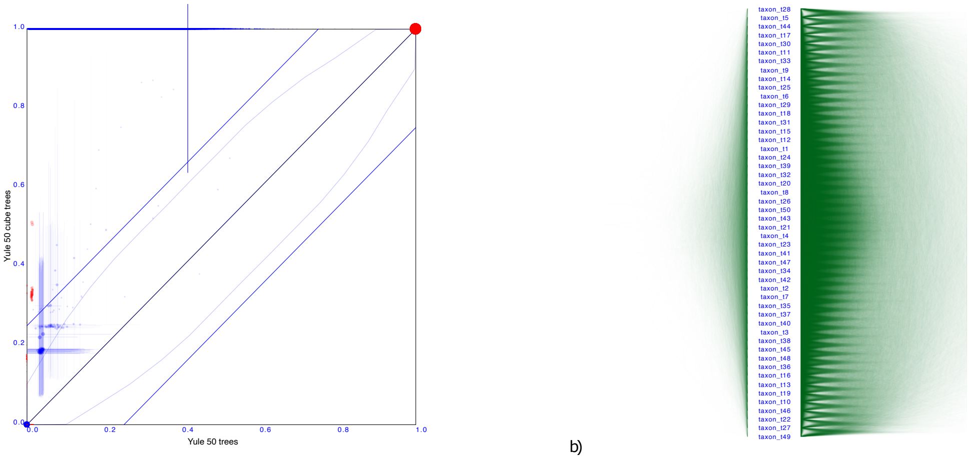


a) b)

**Figure S1** As Figure 3, but with a tree sampled from a Yule prior with 50 taxa, resulting in a tree set sampled uniformly over ranked trees and has high entropy. The cube representation of this tree set has substantially older internal nodes and higher clade support for many clades than the original tree set.
